# Inhibition of delta-1 glutamate receptor current by extracellular protons

**DOI:** 10.64898/2026.06.30.735595

**Authors:** Andrew G. Kain, Joe P. Deuitch, Ayank Maiti, Stephanie C. Gantz

## Abstract

Decreases in brain pH are associated with numerous neurological and neuropsychiatric conditions yet the molecular mechanisms linking decreased brain pH with these disorders are incomplete. The ionotropic glutamate receptors (iGluRs) mediate the majority of excitatory neurotransmission in the brain and can be inhibited or potentiated by extracellular protons depending on receptor subtype. The effect of protons on the delta-glutamate receptor subclass of iGluRs is unknown. Using whole-cell patch-clamp recordings of serotonin neurons in mouse brain slices and activating alpha 1-adrenergic receptors to induce delta-1 glutamate receptor (GluD1_R_) current, we demonstrated that GluD1_R_ current is reduced by physiological drops in extracellular pH. Using two-electrode voltage-clamp recordings in *X. Laevis* oocytes, we further demonstrated that current from a constitutively active variant, GluD1_R_-A650T, was also reduced by extracellular protons. Finally, mice lacking GluD1_R_ showed impaired behavioral responses to inhalation of CO_2_. Taken together, this study continues to expand on the growing body of evidence positing GluD1_R_ as a functional ion channel and suggests that GluD1_R_ facilitates pH sensing *in vivo*.

## Introduction

Small deviations (0.05 units) from neutral blood pH (~7.4) have profoundly negative consequences to brain health [1,2]. These deviations can arise from numerous causes including stroke, heart attack, seizures, head trauma, pneumonia and other chronic lung diseases, obstructive sleep apnea, pain, stress, and anxiety-related hyperventilation can cause long-lasting cognitive impairment. Recently, it has been proposed that decreases in brain pH contribute to cognitive decline in aging, which is exacerbated in Alzheimer’s disease [3], and neuropsychiatric symptoms in bipolar disorder and schizophrenia [4]; however, the molecular mechanisms that sense physiologically relevant shifts in pH and create long-lasting alterations in neuronal excitability are incomplete. For example, non-inactivating voltage-gated potassium channels carry persistent current but are only affected at very low pH (<6.5) [5]. Acid-sensing ion channels (ASICs) are activated by physiologically relevant drops in pH but rapidly desensitize [6–8] often carrying only a small, sustained current [9]. Thus, there is a critical need to identify the primary mode by which neurons convert a change in extracellular pH to a change in the excitability of the neural network over long periods of time. Such a characterization may reveal novel, druggable targets to mitigate the negative consequences of a decrease in brain pH, irrespective of the cause.

The ionotropic glutamate receptors (iGluRs) are all sensitive to extracellular protons in diverse ways. Current carried by the majority of AMPA_R_, NMDA_R_, and kainate_R_ subtypes is reduced by extracellular protons with average IC_50_ values of approximately 6.2, 7.3, and 7.4, respectively [10–12]. Current carried by other subtypes, including GluN1/GluN3A NMDA_R_ and GluK2/GluK4 kainate_R_ is potentiated by extracellular protons [12–14]. Far less is known about the delta glutamate receptors (GluD1_R_ and GluD2_R_). The proton sensitivity of GluD_R_ has only been examined using a constitutively open-pore variant GluD2_R_ ‘*Lurcher*’ which appears similar to NMDA_R_ [15]. This is largely due to the historic difficulty in studying the ion channel function of GluD_R_. However, numerous studies using acute mouse brain slices have shown that GluD_R_ are functional ion channels that can be modulated by G protein-coupled receptor (GPCR) activity [16–19]. In 2020, we discovered that synaptic activation of G protein-coupled *α*1-adrenergic receptors in dorsal raphe serotonin neurons produces a long-lasting excitatory postsynaptic current via coupling to GluD1_R_ channels (*α*1-A_R_-GluD1_R_ current) [18,20]. We and others have reported that GluD1_R_ are open at a low-level and carry a reproducible, excitatory ‘tonic’ current under basal recording conditions [18,21,22]. Thus, in native brain tissue, GluD1_R_ carries a persistent excitatory current that alters neuronal excitability over long periods of time. Moreover, human GluD1_R_ variants have been linked to bipolar disorder, schizophrenia, alcohol use disorder, autism spectrum disorder, and major depressive disorder [19]; however, the extent and mechanism by which GluD1_R_ variants contribute to these disorders is largely unknown.

Here, using whole-cell patch-clamp electrophysiology, we found that synaptic *α*1-A_R_-GluD1_R_ current and tonic GluD1_R_ current were reduced by extracellular protons. Further in *X. Laevis* oocytes, current from a constitutively active variant, GluD1_R_-A650T, was reduced by extracellular protons. As physiological drops in extracellular pH reduce GluD1_R_ current, we hypothesized that GluD1_R_ may function *in vivo* as a pH chemosensor. Indeed, global deletion of GluD1_R_ in mice reduced the behavioral response to elevated levels of CO_2_.

## Methods

### Animals

All experiments were conducted in accordance with the University of Iowa with the approval of the University of Iowa Institutional Animal Care and Use Committee. All electrophysiology experiments used male and female C57BL/6J obtained from The Jackson Laboratory (>2 months old, #000664). All behavioral assays used male and female GluD1^+/+^ and GluD1^-/-^ mice (mixed 129/SvEv and C57BL/J strain) originally generated by Dr. Jian Zuo (St. Jude Children’s Research Hospital). Mice were group-housed on a 12/12 light/dark cycle.

### Acute brain slice preparation and electrophysiological recordings

Brain slices were made as previously described [18,20,21,23]. In brief, mice were anesthetized with isoflurane and euthanized via rapid decapitation. Brains were quickly removed and placed in warmed modified Krebs’ buffer (bubbled with 5/95% CO_2_/O_2_) containing (in mM): 126 NaCl, 2.5 KCl, 1.2 MgCl_2_, 1.2 CaCl_2_, 1.2 NaH_2_PO_4_, 21.5 NaHCO_3_, 11 D-glucose and 5 µM MK-801 to reduce excitotoxicity during slice collection. Coronal slices (240 µm) were collected with a vibrating microtome (Leica) and incubated in the modified Krebs’ buffer at 28 °C for at least 30 min before recording.

Brain slice electrophysiological recordings were made from serotonin neurons in the dorsal raphe nucleus, as previously described [18,20,21,23]. Slices were continuously perfused with modified Krebs’ buffer maintained at 35 °C with an in-line heating element and temperature controller (Warner Instruments). Recordings were made with MultiClamp 700B amplifiers, Digidata 1440A and 1550B converters, and ClampEx software (Molecular Devices, ver. 10.7 and 11.3) with borosilicate glass electrodes (World Precision Instruments and Warner Instruments) wrapped in parafilm to minimize pipette capacitance. Series resistance was monitored throughout the recordings. All reported voltages are corrected for an −8-mV liquid junction potential between the internal and external solutions. α1-A_R_-excitatory postsynaptic currents (α1-A_R_-GluD1_R_ current) (V_hold_ = −65 mV) were evoked every 90-s by trains of current (5 pulses, 0.5 ms, 60 Hz) delivered to the brain slice via a glass monopolar stimulating electrode filled with modified Krebs’ buffer. The stimulating electrode was placed within 200 µm of the recorded neuron. All drugs were applied by bath application, and pH was reduced by bubbling the modified Krebs’ buffer with 20/80% CO_2_/O_2_ gas mixture for at least 30 min at 35 °C prior to application. All recordings were made in the presence of GluN (MK-801, 5 μM), GluA/GluK (NBQX, 3 µM), GABA_A_ (picrotoxin, 100 µM), and 5-HT1A (WAY-100635, 300 nM) receptor antagonists to isolate α1-A_R_-GluD1_R_ current. For complete methodology, see supplementary materials and methods.

### Fast-Scan Cyclic Voltammetry (FSCV)

FSCV was used to measure the rate of pH change when switching from pH 7.4 to 6.8 Kreb’s buffer following previously described methods [24]. For complete methodology, see supplementary materials and methods.

### Stationary Fluctuation Analysis

Stationary fluctuation analysis was conducted as previously described [18,21]. In brief, a single recording in pH_e_ 7.4 and 6.8 were selected for each cell, and the baseline variance was subtracted from the variance at the peak of the α1-A_R_-GluD1_R_ current for each recording. The membrane variance (σ^2^) divided by the macroscopic current (*I*) gives estimates of unitary current (*i*) and open probability (*P*_o_) via:

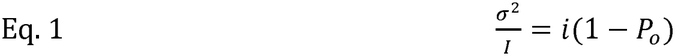

The σ^2^-I relationship was well-fit with a simple linear regression [18], suggesting that P_o_<<0.5 (see [25]) and Eq. 1 can be simplified to:

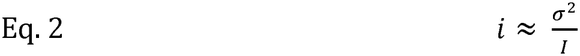

To calculate single-channel chord conductance (γ), we used the following relationship:

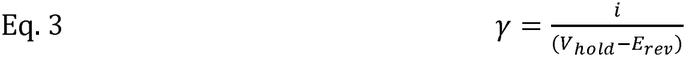

Where V_hold_ is the holding voltage (−65 mV), and E_rev_ is the reversal potential of the α1-A_R_-GluD1_R_ current obtained from the voltage-ramp experiments. Single-channel conductance was calculated with the corresponding E_rev_ obtained in pH_e_ 7.4 and 6.8.

### Molecular biology and two-electrode voltage-clamp recordings in X. Laevis oocytes

Human GluD1_R_ (*GRID1*, NM_017551) DNA of the constitutively active A650T point mutation [26] *GRID1* gene fragments were synthesized by Integrated DNA Technologies, amplified with standard PCR and inserted into the pcDNA3.1 vector via NEBuilder® HiFi DNA Assembly Master Mix (New England Biolabs). The full plasmid sequence was verified with whole-plasmid sequencing (plasmidsaurus).

*Xenopus laevis* oocytes were prepared from ovaries (Xenopus 1, Inc). Healthy stage V and VI defolliculated oocytes were manually selected and stored in modified OR3 media at 17 °C. The next day, individual oocytes were injected with ~36 ng of RNA with a nanoliter 2020 microinjector (World Precision Instruments). Injected oocytes were maintained in OR3 media for 24 h before electrophysiology experiments were conducted.

Two-electrode voltage-clamp required were acquired at 10 kHz using a Warner OC-725C amplifier, Digidata 1440A digitizer, and ClampEx software (Molecular Devices, ver. 10.7). Pipette resistances were ≤0.4 MΩ when filled with 3M KCl. Oocytes were placed in the Xenoplace Workstation bath (ALA), and gravity-perfused with Na^+^-Ringer’s solution containing (in mM): 90 Na^+^, 3 KCl, 0.5 BaCl_2_, 10 HEPES. pH was adjusted to 6.0, 7.0, or 8.0 with NaOH prior to adding NaCl to ensure each solution had 90 mM Na^+^. NMDG-Ringer’s solution at pH 8.0 was used and contained (in mM): 90 NMDG^+^, 3 KCl, 0.5 BaCl_2_, 10 HEPES, pH adjusted with HCl. Constitutive inward current was measured as described previously [26] from V_hold_ of −35 mV. For complete methodology, see supplementary materials and methods.

### Behavioral assays

For all behavioral experiments, mice were placed in a rabbit CO_2_ induction chamber (18”L× 12”W×12”H; EZ-179; EZ Systems) fit with a CO_2_ sensor (USB-DXC220t-CAL; Dracal Technologies). Mice habituated to the chamber for 15 minutes on preceding days before testing on day 3. During habituation and air exposure, room air circulated through the chamber using an aquarium pump. To expose mice to elevated levels of CO_2_, the aquarium pump line was clipped off and a line connecting a gas cylinder (20% CO_2_, 21% O_2_, N_2_ balance) was opened. The CO_2_ gas mix was pumped into the chamber for 10 min at a rate of 5.0 L/min resulting in a maximum level of CO_2_ in the chamber of ~8.5%. Higher flow rates were not used as they resulted in an increase in sound intensity in the chamber. Trials were recorded using a camera mounted above the CO_2_ chamber. Total time spent freezing was scored manually from these videos using the free Behavioral Observation Research Interactive Software (BORIS), with the scoring investigator blind to the sex, genotype, and condition.

### Experimental design and statistical analysis

All data were analyzed using ClampFit 10.7 and 11.1. Data are presented as representative current traces, or as means±S.E.M, with each point representative of a unique cell. ‘N’ denotes the number of cells for all electrophysiological experiments. All data sets are analyzed with nonparametric statistical tests (Wilcoxon matched-pairs signed rank test for two-group within analyses, Mann-Whitney tests for two-group between analyses, Kruskal-Wallis test for three or more between-group analyses, Friedman test - or mixed-effects analysis if conditions had unequal ‘n’s’ - for three or more within-group analyses, or 2-way ANOVA). All datasets were checked for sex differences, and statistically significant sex differences are reported. An alpha value of 0.05 was used to determine statistical significance, and exact p-values are reported unless p<0.001 or >0.999. All statistical analyses were performed in GraphPad Prism (GraphPad Software, Inc.).

## Results

### *α*1-A_R_-GluD1_R_ current is reduced by extracellular protons

Whole-cell voltage-clamp (V_hold_ −65 mV) electrophysiological recordings were obtained from serotonin neurons in the dorsal raphe nucleus in acute mouse brain slices at 35*°* C using a potassium-based internal solution with NMDA_R_, AMPA_R_ and Kainate_R_, GABA-A_R_ and 5-HT1A_R_ antagonists in the external solution. A train of electrical stimuli (5 pulses, 60 Hz, every 90s) was used to reliably evoke synaptic *α*1-A_R_-GluD1_R_ current, as previously described [18,20,21,23]. The pH of the extracellular (pH_e_) bicarbonate-buffered artificial cerebral spinal fluid (aCSF) was changed from the standard 7.4 to 6.8 by increasing the concentration of saturated CO_2_ from 5% to 20%. Application of pH_e_ 6.8 reduced *α*1-A_R_-GluD1_R_ current from −52±7 pA to −21±3 pA and was readily reversed (Fig.1A-C). To compare the rates of reduction and recovery of *α*1-A_R_-GluD1_R_ current to the real-time change in pH_e_, pH_e_ of the bulk solution was measured using carbon fiber fast-scan cyclic voltammetry [24]. The reduction of *α*1-A_R_-GluD1_R_ current was well-aligned with the decrease in pH_e_, whereas the return to pH_e_ 7.4 preceded full recovery of *α*1-A_R_-GluD1_R_ current by ~4.5 minutes (Fig.1C). *α*1-A_R_-GluD1_R_ current kinetics were also changed significantly by pH_e_ 6.8, with the time constant of activation (*τ*-activation) increasing from 359±37 ms in pH_e_ 7.4 to 771±135 ms in pH_e_ 6.8 (Fig.1D-E). The time constant of decay (*τ*-decay) was inversely affected by protons, decreasing from 7.92±0.87 s in pH_e_ 7.4 to 6.17±0.72 s in pH_e_ 6.8 (Fig.1F). In a prior report, we demonstrated that the amplitude of *α*1-A_R_-GluD1_R_ current does not affect the rates of activation and decay [20], so it is unlikely the reduction in amplitude by pH_e_ 6.8 underlies the change in time course.

**Fig. 1.**
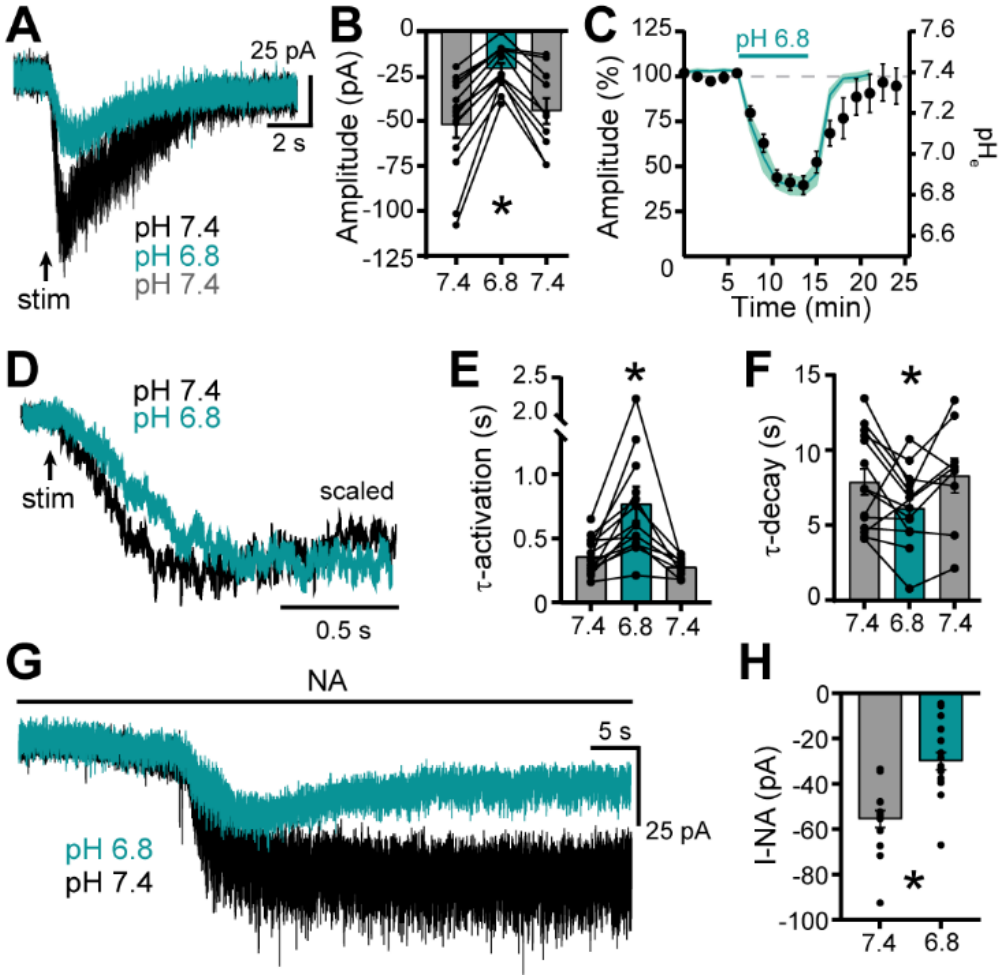
Extracellular protons reduce the *α*1-A_R_-GluD1_R_ current. **A** Representative traces showing the *α*1-A_R_-GluD1_R_ current in pH_e_ of 7.4 (black), 6,8 (teal), and recovery in 7.4 (gray). **B** Quantification of the *α*1-A_R_-GluD1_R_ current amplitude while acidifying the bath recording solution and recovery upon returning to 7.4 pH_e_ (7.4 v 6.8, p<0.001, n=14; 7.4 v 7.4, p=0.379, n=10, REML with Holm-Šídák’s multiple comparisons test). **C** Time series of the *α*1-A_R_-GluD1_R_ current amplitude during bath acidification and wash back to pH_e_ 7.4 (n=10-14). The teal line is the quantification of the real-time change in bath pH measured with FSCV (n=3). **D** Representative traces showing the activation kinetics of the *α*1-A_R_-GluD1_R_ current at pH_e_ 7.4 (black) and 6.8 (teal). **E** Quantification of the activation kinetics of the *α*1-A_R_-GluD1_R_ current at pH_e_ 7.4, 6.8, and recovery back in 7.4 (7.4 v 6.8, p=0.006, n=14; 7.4 v 7.4, p=0.168, n=9, REML with Holm-Šídák’s multiple comparisons test). **F** Quantification of the decay kinetics of the *α*1-A_R_-GluD1_R_ current at pH_e_ 7.4, 6.8 and recovery back in 7.4 (7.4 v 6.8, p=0.046, n=13; 7.4 v 7.4, p=0.635, n=9, REML with Holm-Šídák’s multiple comparisons test). **G** Representative traces showing the current elicited by bath application of 30 μM noradrenaline (I-NA) at pH_e_ 7.4 (black) and 6.8 (teal). (**H**) Quantification of the I-NA amplitude at pH_e_ 7.4 and 6.8 (p<0.001, n=15 and n=17, respectively, Mann-Whitney test).

Decreasing extracellular pH can reduce neurotransmitter release via inhibition of voltage-gated calcium channels [27]. To determine whether extracellular protons reduced *α*1-A_R_-GluD1_R_ current via presynaptic inhibition of noradrenaline release, we bypassed the presynaptic element and bath-applied noradrenaline onto the brain slice (30 *µ*M, in the presence of 300 nM idazoxan [23]). In pH_e_ 7.4, noradrenaline produced an average inward current of −55±4 pA, consistent with prior reports [18,20] (I-NA, Fig.1G-H). Decreasing pH_e_ to 6.8 reduced the average amplitude of I-NA by ~45% to −30±4 pA (Fig.1G-H), indicating that proton-mediated reduction of *α*1-A_R_-GluD1_R_ current is not due to presynaptic mechanisms. Overall, these results show that extracellular acidification by −0.6 units from neutral pH reduces *α*1-A_R_-GluD1_R_ current via a postsynaptic mechanism.

### Protons reduce *α*1-A_R_-GluD1_R_ unitary current

With the exception of GluN1/GluN3A NMDA_R_ and GluK2/GluK4 Kainate_R_, current carried by iGluRs is reduced by extracellular protons in a voltage-insensitive manner that largely reduces the open probability and mean open time without altering channel conductance [10–13,28–31]. However, recent evidence suggests a slight decrease in calcium permeability for NMDA_R_ [32]. To determine how protons reduce *α*1-A_R_-GluD1_R_ current, we examined the current-voltage relationship generated by voltage ramps (−120 mV to +30 mV, 1 mV/10 ms) during the *α*1-A_R_-GluD1_R_ current in pH_e_ 7.4 and 6.8 (Fig.2A). Consistent with our prior report [18], the current reversed polarity at −32.0±3.2 mV in pH_e_ 7.4. In pH_e_ 6.8, the current was reduced across the voltage range, without shifting the reversal potential (−29.8±3.1 mV, Fig.2B). The effect of protons was voltage-insensitive (Fig.2C), suggesting action at a site distal to the channel pore.

A feature of *α*1-A_R_-GluD1_R_ current is a significant increase in membrane noise variance (*σ*^2^) [18] (Fig.2D). Using between-cell stationary fluctuation analysis (with the assumption that the macroscopic current and *σ*^2^ arise from identical, independent channels opening and closing), single-channel properties of GluD1_R_ were estimated [25,33]. The *α*1-A_R_-GluD1_R_ current *σ*^2^-amplitude relationship in pH_e_ 7.4 and 6.8 was well-fit by a linear regression [18,21] (Fig.2E), suggesting that open probability is small (<<0.5). In pH_e_ 7.4, mean unitary current was ~-1.12±0.1 pA, which was reduced by ~45% in pH_e_ 6.8 to ~-0.62±0.1 pA (Fig. 2F). Single-channel chord conductance was reduced by ~45% from 34.1±3.1 pS in pH_e_ 7.4 to 18.7±3.4 pS in pH_e_ 6.8 (Fig.2G).

**Fig. 2.**
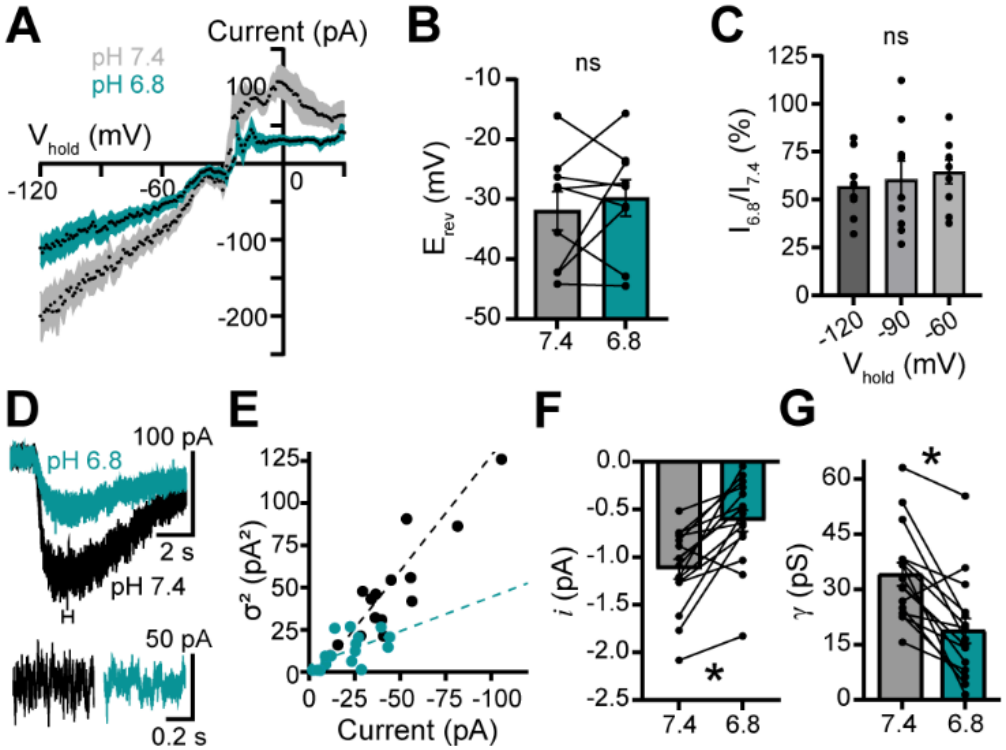
Protons reduce *α*1-A_R_-GluD1_R_ current by reducing unitary current. **A** Background-subtracted voltage ramps (−120 mV to +30 mV, 1 mV/10 ms) collected at the peak of the *α*1-A_R_-GluD1_R_ current at pH_e_ 7.4 (gray, n=9) and 6.8 (teal, n=9). The shaded area represents the S.E.M. **B** Quantification of the reversal potential of the *α*1-A_R_-GluD1_R_ current at pH 7.4 and 6.8 (p=0.734, n=9, Wilcoxon matched-pairs signed rank test). **C** Quantification of the voltage-dependence of proton-mediated inhibition. For each holding potential, the current observed ay pH 6.8 was divided by the current seen at pH 7.4 and converted to a percentage (p=0.814, n=9, Friedman test). **D** Representative traces of the peak noise of the *α*1-A_R_-GluD1_R_ current at pH_e_ 7.4 (black) and 6.8 (teal). Noise was measured in a 700 ms window at the peak of the *α*1-A_R_-GluD1_R_ current. **E** Stationary fluctuation analysis measured at the peak of the *α*1-A_R_-GluD1_R_ current at pH_e_ 7.4 (black, n=16) and 6.8 (teal, n=16). The data was well-fit with a linear regression. **F** Quantification of mean GluD1_R_ unitary current (**i**). Unitary current was calculated for each cell via Eq.2 (p<0.001, n=16, Wilcoxon matched-pairs signed rank test). **G** Quantification of single-channel chord conductance using Eq.3 and the respective E_rev_ measurements obtained in **B** (p<0.001, n=16, Wilcoxon matched-pairs signed rank test).

### Tonic GluD1_R_ current is reduced by extracellular protons

In cell lines and brain slices, GluD1_R_ and GluD2_R_ are open and carry tonic cation current [18,21,22,34]. While the origin of tonic GluD1_R_ current remains unknown, it is not affected by arresting GTP/GDP exchange nor elevating free G*βγ* subunits [21], allowing for measurements of GluD1_R_ channel activity separate from G protein-mediated GluD1_R_ currents. Tonic GluD1_R_ current was revealed by the open-channel pore blocker 1-napthyl acetyl spermine (NSP, 100 *µ*M, Fig.3A), which produced an apparent outward current at pH_e_ 7.4 and 6.8 (Fig.3A-B). When compared with measurements in pH_e_ 7.4, the magnitude of tonic GluD1_R_ current in pH_e_ 6.8 was reduced by ~46% from −32.6±4.7 pA to −17.6±2.9 pA in pH_e_ 6.8 (Fig.3C). A feature of the tonic GluD1_R_ current is a significant decrease in membrane noise variance (*σ*^2^) upon NSP application [18,21]. On average in pH_e_ 7.4, NSP reduced *σ*^2^ from 17.8±1.9 pA^2^ to 13.8±2.0 pA^2^ (Fig.3A,D). In contrast, in pH_e_ 6.8, *σ*^2^ was 13.9±3.0 pA^2^ and was not further reduced by NSP (13.0±2.6 pA^2^) (Fig.3A,D).

**Fig. 3.**
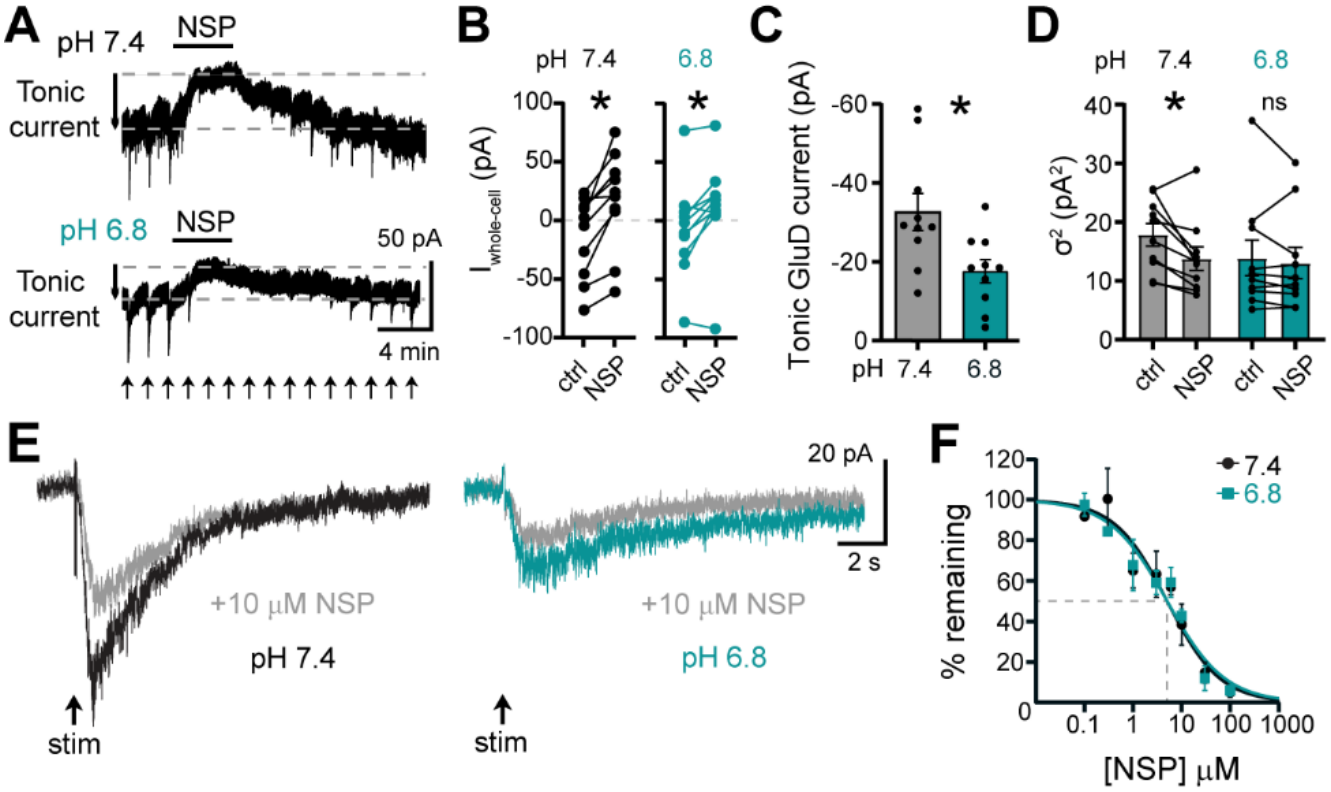
Tonic GluD1_R_ current is reduced by extracellular protons. **A** Representative traces showing tonic GluD1_R_ current as revealed by application of 100 µM NASPM (NSP) in pH_e_ 7.4 (top) and 6.8 (bottom). **B** Quantification of whole-cell current immediately preceding (control) and at the peak of the NASPM current (NSP) at pH_e_ of 7.4 (black, p=0.002, n=10, Wilcoxon matched-pairs signed rank test) and 6.8 (teal, p=0.020, n=10, Wilcoxon matched-pairs signed rank test). **C** Quantification of the tonic GluD1_R_ current peak amplitude in pH_e_ 7.4 (gray) and 6.8 (teal, p=0.015, n=10, Mann-Whitney test). **D** Quantification of the change in membrane noise variance (σ^2^) caused by 100 μM NASPM at pH_e_ 7.4 (grey) and pH_e_ 6.8 (teal). Measurements of σ^2^ were obtained in a 700 ms window (7.4, p=0.004, n=10; 6.8, p=0.496, n=10, 2-way ANOVA with Fischer’s LSD test). **E** Representative traces showing the *α*1-A_R_-GluD1_R_ current inhibition by a submaximal (10 *µ*M) application of NASPM in pH_e_ 7.4 (black, left) and 6.8 (teal, right). **F** Concentration-response curve of the *α*1-A_R_-GluD1_R_ current to varying concentrations of NASPM at pH_e_ 7.4 (black) and 6.8 (teal), demonstrating equivalent steady-state potency and efficacy (IC_50_ 7.4: 5.35 μM, 6.8: 4.27 μM, p=0.544).

In principle, a decrease in tonic GluD1_R_ current could be explained if the ability of NSP to block GluD1_R_ channels was impaired. Thus, we generated a concentration response curve to determine the efficacy and potency of NSP to block *α*1-A_R_-GluD1_R_ current (Fig.3E-F). NSP (100 *µ*M) nearly eliminated *α*1-A_R_-GluD1_R_ current in both pH_e_ 7.4 and pH_e_ 6.8, demonstrating equivalent efficacy (Fig.3F). In pH_e_ 7.4 and pH_e_ 6.8, the concentration of NSP required to achieve 50% block (IC_50_) was not statistically different; 5.35 *µ*M and 4.27 *µ*M, respectively (Fig.3E-F). Taken together, the results show that extracellular protons reduce tonic GluD1_R_ current, suggesting direct action on GluD1_R_ channels.

### Protons reduce constitutively active GluD1_R_-A650T current in *X. Laevis oocytes*

*Het*erologous expression systems are instrumental in studying the biophysical properties of numerous ion channels as they provide a simplified model system. Since current from wild type GluD_R_s has not been identified in heterologous expression systems, the use of constitutively active variants has been vital towards advancing the study of these enigmatic receptors [15,35]. Here, two-electrode voltage-clamp recordings were made from *Xenopus Laevis* oocytes expressing a human GRID variant, GluD1_R_-A650T, which produced a constitutive inward current in the presumed absence of ligand [36] (Fig.4). Consistent with a prior report [26], D-serine (10 mM) potentiated GluD1_R_-A650T current by ~40% (Fig.4A-B). Next, the effect of protons on GluD1_R_-A650T was examined. As the concentration of protons was reduced from pH_e_ 6.0 to 7.0 and 8.0, there was significant potentiation of GluD1_R_-A650T (Fig.4A-B), consistent with a prior report on the constitutively active GluD2_R_-A654T ‘*Lurcher*’ variant [15]. Taken together, these results indicate that when expressed in a heterologous expression system, GluD1_R_ current is reduced directly by extracellular protons.

**Fig. 4.**
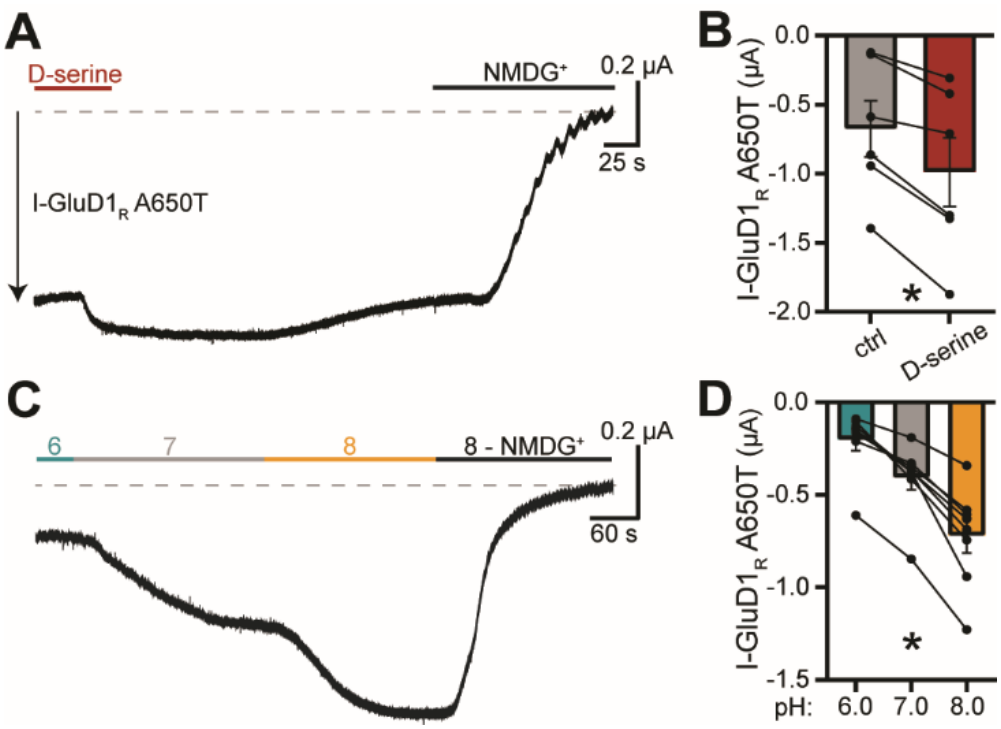
Current carried by the constitutively active GluD1_R_-A650T variant is reduced by extracellular protons in *X. Laevis oocytes.* ***A*** Representative trace showing potentiation of GluD1_R_-A650T current by 10 mM D-serine. At the end, NMDG^+^ was applied to facilitate measurement of the constitutive inward current (I-GluD1_R_-A650T). **B** Quantification of 10 mM D-serine potentiation of GluD1_R_-A650T current (p=0.031, n=6, Wilcoxon matched-pairs signed rank test). **C** Representative trace showing GluD1_R_-A650T current is potentiated as the extracellular proton concentration is reduced from pH 6.0 to pH 7.0 to pH 8.0. At the end, NMDG^+^ was applied to facilitate measurement of the constitutive inward current (I-GluD1_R_-A650T). **D** Quantification of the pH sensitivity of GluD1_R_-A650T. Reducing the concentration of extracellular protons potentiated GluD1_R_-A650T current (p<0.001, n=8, Friedman test; 6.0 v 7.0: p=0.046; 7.0 v 8.0: p=0.046, Uncorrected Dunn’s test).

### Mice lacking GluD1_**R**_ **have reduced sensitivity to elevated levels of CO**_**2**_

To determine whether GluD1_R_ functions as a pH chemosensor *in vivo*, wild type or global GluD1_R_ knockout mice (GluD1_R_-KO) inhaled elevated levels of CO_2_. In mice, inhalation of 10% CO_2_ reduces brain pH from 7.15 to 6.97 [37]. In brief, an induction chamber was equipped with an inlet to deliver room air or CO_2_, an outlet for CO_2_ scavenging, and a CO_2_ sensor. Mice habituated to the chamber for 15 minutes for two consecutive days before being exposed to either air or 8.5% CO_2_ on the third day (Fig.5A). Air or CO_2_ was administered to the chamber for 10 minutes starting after the mouse was placed inside and the chamber sealed. 90 minutes after the end of the trial, mice were perfused and their brains collected. When compared to wild type mice, GluD1_R_-KO mice froze significantly less over the 15-minute exposure to CO_2_ (Fig.5B,D). As a metric of the behavioral sensitivity to CO_2_, the CO_2_ level when each mouse had spent a cumulative time freezing of 1.5 mins (10% of the trial) was recorded. On average, wild type mice froze for 10% of the trial at 4.8±0.4% CO_2_ (Fig.5C). In contrast, GluD1_R_-KO mice required 6.1±0.4% CO_2_ (Fig.5C). Overall, these results suggest that mice lacking GluD1_R_ are less sensitive to elevated levels of CO_2_, positing GluD1_R_ as an *in vivo* pH-sensor.

**Fig. 5.**
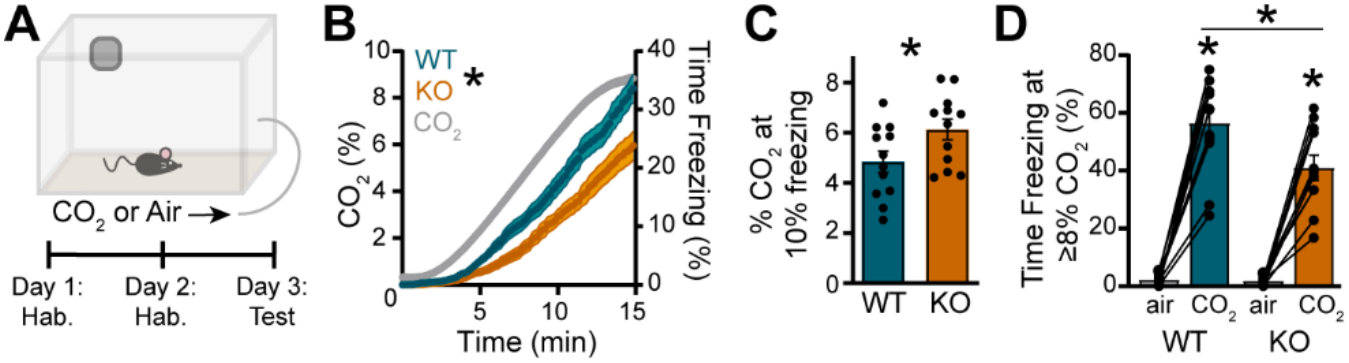
Mice lacking GluD1_R_ display a reduced sensitivity to elevated levels of CO_2_. **A** Cartoon schematic detailing the behavioral paradigm. Air was administered on days 1 and 2. Either CO_2_ or air was administered on day 3. **B** Time series of cumulative time freezing (right axis) for GluD1-WT (blue) and GluD1-KO (orange) mice during CO_2_ exposure (gray line, left axis). CO_2_ was administered for 10 of the 15-minutes trial before being turned off, resulting in a consistent steady state value of ~8.5% (n=24). **C** Quantification of the percentage of CO_2_ in the testing chamber that elicited 10% of the total time spent freezing for GluD1-WT (blue) and GluD1-KO (orange) mice (p=0.040, n=12, Welch’s t-test). **D** Quantification of the cumulative time spent freezing for GluD1-WT (blue) and GluD1-KO (orange) mice (WT, p<0.001, n=12; KO, p<0.001, n=11; WT v KO in CO_2_, p=0.002, n=11-12, 2-way ANOVA with Fischer’s LSD test).

Towards determining whether one region or cell-type may underly the behavioral phenotype observed in the global GluD1_R_-KO mice, GluD1_R_-dependent differences in neuronal activation driven by CO_2_ exposure were assayed using immunostaining of the early gene product, c-Fos [38] in brain regions implicated in freezing and arousal behaviors to CO_2_, including the dorsal raphe [39], locus coeruleus (LC) [40], and central amygdala (CeA) [6,41,42] which all express GluD1_R_ [43]. In the dorsal raphe, c-Fos^+^ serotonin neurons were scarce (<~5%) and there were no differences between conditions nor genotypes (Fig.S2A-C). In the LC, exposure to CO_2_ caused a significant increase in c-Fos^+^ noradrenergic neurons comparably between wild type and GluD1_R_-KO mice (Fig.S2D-F). In the CeA, exposure to CO_2_ caused a significant increase in c-Fos^+^ PKCδ^+^ neurons, from GluD1_R_-KO but not wild type mice (Fig.S3). These data suggest that loss of GluD1_R_ reduces freezing responses to elevated CO_2_, at least in part via increased activation of CeA PKCδ^+^ neurons.

## Discussion

### Mechanism of proton reduction of GluD1_R_ current

Proton sensitivity of the constitutively open GluD2_R_-A654T ‘*Lurcher*’ variant suggested that native GluD_R_ may also be reduced by extracellular protons [15]. Here, we found that extracellular protons in a physiologically relevant range (pH_e_ 6.8) reduce native GluD1_R_ current significantly. The reduction was not dependent on presynaptic mechanisms, and the magnitude of reduction for both *α*1-A_R_-GluD1_R_ current and tonic GluD1_R_ current was similar. Further, constitutively open GluD1_R_-A650T current recorded in *X. Laevis* oocytes was also reduced by extracellular protons, suggesting that protons exert direct action on GluD1_R_ channels.

The mechanism by which protons reduce macroscopic GluD1_R_ current is not well-resolved. Extracellular protons largely reduce NMDA_R_, AMPA_R_, and kainate_R_ current with some exceptions being potentiated [10–15]. When iGluR currents are reduced by protons, protons reduce the probability of channel opening and mean open time without altering channel conductance [10–12,28–31]. Here, using stationary noise analysis of macroscopic GluD1_R_ current, we observed a proton-mediated decrease in the estimate of GluD1_R_ conductance, without a significant change the relative permeability of cations which is expected to shift the current-voltage relationship. However, since the *σ*^2^-current amplitude relationship was well-fit with a linear regression, instead of a parabolic relationship [25], and membrane noise was measured over hundreds of milliseconds, these analyses are unable to accurately define the probability of opening (P_o_). Thus, a reduction in P_o_ may contribute to the observed proton-mediated decrease in the estimate of GluD1_R_ conductance. Indeed, P_o_ and mean open time shapes the time course of activation and decay of macroscopic currents [44]. Proton-mediated reductions in P_o_ and mean open time of GluN1/GluN2B NMDA_R_ and GluA2 AMPA_R_ are accompanied by increases in the τ-activation and decreases in τ-decay [30, 45], similar to the changes we observed in τ-activation and τ-decay of the *α*1-A_R_-GluD1_R_ current in pH_e_ 6.8. These data suggest that protons may also reduce the probability of opening and/or mean open time of GluD1_R_. Ultimately, single-channel recordings of GluD1_R_ current and resolving the structure of GluD1_R_ under acidic conditions will be necessary to define the precise mechanism(s).

### Health implications

Acidic shifts in brain pH can arise from cardiovascular, pulmonary, neurological, and neuropsychiatric disorders and can cause cognitive impairment. In this study, we demonstrate that long-lasting excitatory current carried by GluD1_R_ channels is reduced by extracellular protons. *In vivo*, noradrenergic activation of α1-A_R_ drives serotonin neurons to fire action potentials [46] and release serotonin [47,48] via GluD1_R_ conductance [18]. Thus, a decrease in GluD1_R_ current in serotonin neurons, in principle, would promote a decrease in serotonin neuronal excitability with a decrease in brain pH. Reduced excitability of serotonin neurons and subsequent serotonin release is expected to alter mood and affect, sleep/wake control, and thermoregulation, which may contribute to the symptoms of depression, anxiety, bipolar disorder, and schizophrenia. In humans, variations in the gene encoding GluD1_R_ (GRID1) have been linked to many of the same disorders [19,26] although how these genetic variations impact the ion channel function of GluD1_R_ is largely unknown. One exception is the schizophrenia-associated constitutively open GluD1_R_-A650T variant [26], which we found retains proton-sensitivity but still carries inward current at pH_e_ 6.0. Future work is necessary to determine how GluD1_R_-A650T contributes to the pathophysiology of schizophrenia.

In the present study, global GluD1_R_ knock-out mice displayed reduced freezing responses to elevated levels of CO_2_, which was accompanied by increased activity of CeA PKCδ^+^ neurons, only in the GluD1_R_-KO mice. The CeA is a key brain region associated with sensing and driving freezing responses to CO_2_ [6,41,42], at least in part due to activation of CeA neurons via ASICs [6]. Although it should be noted that within the CeA, there are opposing roles of PKCδ^+^ neurons inhibiting and somatostatin^+^ neurons promoting freezing behaviors [42,49] and GluD1_R_ is highly enriched in PKCδ^+^ neurons [50]. Since exposure to elevated CO_2_ had no net effect on the activity of CeA neurons in wild type mice, it is possible that depolarization due to pH-dependent ASIC activation is countered by a pH-dependent loss of excitatory drive via GluD1_R_. With genetic deletion of GluD1_R_, pH-dependent ASIC activation may prevail, causing activation of PKCδ^+^ neurons to inhibit freezing.

While future work will be needed to parse the pH-sensing role of GluD1_R_ within specific brain regions and cell types, this study lays the foundation to further investigate the role of GluD1_R_ in pH chemosensing consequences of proton-mediated channel inhibition in neural network excitability and health following acidic insult. Impaired response to CO_2_ may contribute to mortality due to sleep apnea, sudden unexpected death in epilepsy, and sudden infant death syndrome [51]. Conversely, enhanced sensitivity to CO_2_, for example in individuals with bipolar disorder [52], elicits aversive panic-like states. Overall, these results continue to expand on the growing literature supporting ion channel function of GluD1_R_ with similar biophysical properties as other iGluRs. It also begins to contextualize the physiological role GluD1_R_ play in mediating behavioral responses to changes in brain pH.

## Acknowledgements

We would like to extend our sincere gratitude to Dr. Shashank Dravid (Texas A&M University) for providing the GluD1^+/+^ and GluD1^-/-^ mice. We thank the Ahern lab members Drs. Colin Clark and Dr. Lionel Gissot for their assistance with the molecular biology and *X. Laevis* electrophysiology experiments. The Ahern lab is funded by NIGMS R35GM148239.

## Author Contributions

A.G.K. and S.C.G. conceptualized the study. A.G.K., S.C.G., and J.P.D. collected electrophysiological recordings. A.G.K. and S.C.G. analyzed electrophysiological recordings. A.G.K. and A.M. collected and analyzed behavioral data. A.G.K., S.C.G., A.M. and J.P.D collected and analyzed IHC data. A.G.K. and S.C.G. wrote the manuscript. All authors revised the manuscript.

## Data Availability

The datasets generated during and/or analyzed during the current study are available from the corresponding author on reasonable request.

## AI Declaration

The authors have nothing to disclose.

