## Supplemental Materials for "Inhibition of delta-1 glutamate receptor current by extracellular protons"

**Materials and Methods**

*Acute brain slice preparation and electrophysiological recordings*

Pipette resistances were 3.0-4.5 MΩ when filled with an internal solution containing (in mM): 104.56 K-methysulfate, 3.73 KCl, 5.3 NaCl, 4.06 MgCl_2_, 4.06 CaCl_2_, 7.07 HEPES (K), 3.25 BAPTA (4K), 0.26 GTP-Na salt, 4.87 ATP-Na salt, 4.59 creatine phosphate-Na salt, adjusted to ~275 mOsm and maintained at pH 7.2. For voltage-ramp experiments, a cesium-based internal solution was used containing (in mM): 120 CeMeS, 5.3 NaCl, 7.07 HEPES, 3.25 BAPTA (4K), 0.26 Na-GTP salt, 4.87 Na-ATP salt, 4.59 Na-creatine phosphate, 4.06 MgCl_2_, 4.06 CaCl_2_, adjusted to ~275 mOsm and maintained at pH 7.2.

*Fast-Scan Cyclic Voltammetry (FSCV)*

10 µm carbon fiber microelectrodes (Kation Scientific) were prepared in 200-proof ethanol with activated charcoal for 15 min before being attached to a CV-7B headstage (Molecular Devices) with an HLU-type headstage adaptor (Kation Scientific). Recordings were collected at 4 Hz with a scan rate of 40 V/s and background-subtracted from the average voltammogram collected during the first 2 minutes in pH 7.4 Kreb’s buffer. Calibration curves were generated using six HEPES-buffered solutions of pH 6.8, 7.0, 7.2, 7.4, 7.6, and 7.8. The 7.4 curve was used as the baseline and subtracted from the other curves. Both the ‘Q’ peak and ‘QH’ peak were measured and used to determine bath pH [1]. (Supplementary Figure S1).

*Molecular biology and two-electrode voltage clamp (TEVC) recordings in X. Laevis oocytes*

K-12 E. coli bacteria were transfected with the A650T variant and successful incorporation of the cDNA was verified using gel electrophoresis. Purification of cDNA from the bacteria was done with the Monarch® Spin Plasmid Miniprep Kit (New England Biolabs). The purified cDNA was digested overnight and the cRNA was synthesized using the T7 RNA mMESSAGE kit (Invitrogen).

*X. laevis* oocytes were prepared from ovaries purchased from Xenopus 1, Inc. Upon arrival, the ovary was coarsely dissociated using forceps and rinsed thoroughly in Ca^2+^-free Barth’s solution (CBS) containing (in mM): 88 NaCl, 1 KCl, 2.5 NaHCO_3_, 1 MgSO_4_, 5 HEPES, pH adjusted to 7.4 with NaOH. Once the oocytes were washed and in clean CBS, oocytes were defolliculated using collagenase enzyme (Collagenase Type 4, provided by Xenopus 1) with gentle rocking at room temperature for approximately 45 min. Defolliculated oocytes were then thoroughly washed in CBS before being transferred to modified OR3 media (500 ml Leibovitz’s L-15 media into 500 ml d_2_H_2_O) supplemented with 15 mM HEPES, 0.17 mM Amikacin, 1% penicillin/streptomycin, and amphotericin B at 0.275 µg/ml.

*Immunohistochemistry and confocal microscopy*

For immunohistochemical processing, mice were euthanized with a ketamine/xylazine cocktail and transcardially perfused with 5% sucrose, followed by 4% paraformaldehyde. Brains were extracted, post-fixed overnight in 4% PFA at 4°C. Coronal slices (50 μm and 100 μm) were collected on a vibratome (EMS 5000, Electron Microscopy Sciences, Hatfield, Pennsylvania).

*Central Amygdala:*

Slices were placed in a blocking solution containing 1% Bovine Serum Albumin, 5% normal Donkey Serum, and 0.5% Triton-X for 1 h, then washed in PBS, and incubated in rabbit anti-c-Fos primary (1:4000; Cell signaling technology, Danvers, Massachusetts; 4384) at room temperature overnight. Slices were washed, incubated in donkey anti-rabbit Alexa Fluor 488 (1:1000; Jackson Immunoresearch) at room temperature for 2 h, washed, and incubated overnight at 4° C in mouse anti-PKC delta primary (1:500; BD Biosciences, Franklin Lakes, New Jersey; BDB610397) containing the blocking solution. Slices were washed and incubated in donkey-anti-mouse Alexa Flour 594 (1:250; Jackson Immunoresearch) for 1 h at room temperature. After a final wash, slices were mounted, and coverslipped with Fluoromount-G containing DAPI (SouthernBiotech, Birmingham, AL).

*Dorsal Raphe and LC:*

Slices were placed in a blocking solution containing 0.5% Triton-X and 10% normal donkey serum for 3 h, washed, and incubated with rabbit anti-c-Fos primary (1:4000; Cell signaling technology, Danvers, Massachusetts; 4384) at room temperature overnight. Slices were washed, incubated in donkey anti-rabbit Alex Fluor 488 (1:1000; Jackson Immunoresearch) for 2 h at room temperature and washed. Slices containing the dorsal raphe nucleus were incubated overnight at room temperature with goat anti-TPH2 (1:1000; Abcam, Cambridge, UK; AB1210143). Slices containing the Locus Coeruleus were incubated with chicken anti-TH (1:500; Aves, Davis, California; TYH) overnight at room temperature. Slices were washed, incubated in donkey anti-goat or donkey anti-chicken Alexa Fluor 647, respectively, (1:1000, Jackson Immunoresearch) for 2 h at room temperature. Slices were then washed, mounted, and coverslipped with Fluoromount-G containing DAPI (SouthernBiotech, Birmingham, AL).

*Immunohistochemistry Image Analysis:*

Mounted slices were imaged on a Zeiss Axiovert 100 confocal microscope (Carl Zeiss Microscopy, LLC, White Plains, NY) with a 40× oil-dipping objective. Images were processed in ImageJ and c-Fos expression were manually counted through FIJI software by investigators blinded to the sex and treatment group. The total number of c-Fos positive cells per region per mouse were normalized for the total area imaged (c-Fos/mm^2^).

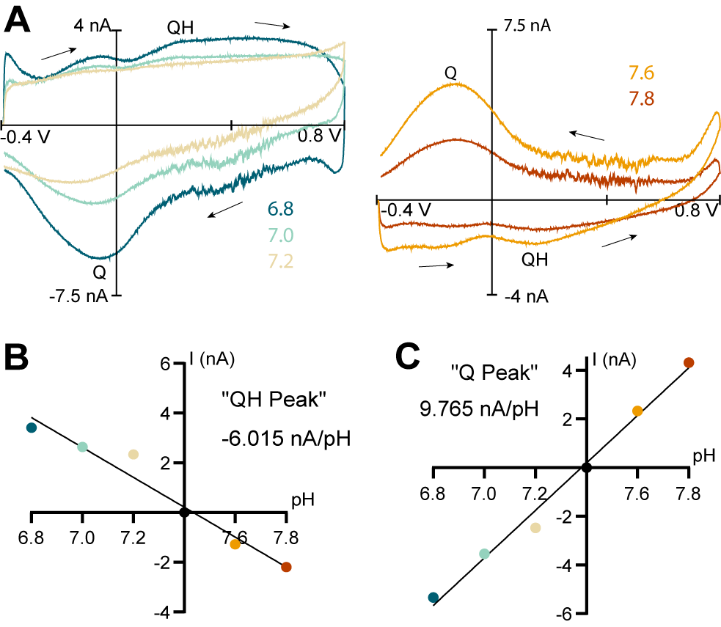

**Supplementary Fig S1**: **Calibration of FSCV curves.**

**A** Representative cyclic voltammograms obtained from -0.4 V to 0.8 V (40 V/s) scans in HEPES-buffered Kreb’s solutions at pH 6.8 (dark teal), 7.0 (light teal), 7.2 (beige), 7.6 (orange) and 7.8 (red). At each pH, the cyclic voltammogram obtained at pH 7.4 was subtracted. Arrows indicate the direction of the scan. **B** Quantification of the QH peak current-pH relationship. **C** Quantification of the Q peak current-pH relationship. Note, both the QH and Q peak values were averaged together to produce the pH timeseries seen in Fig. 1C.

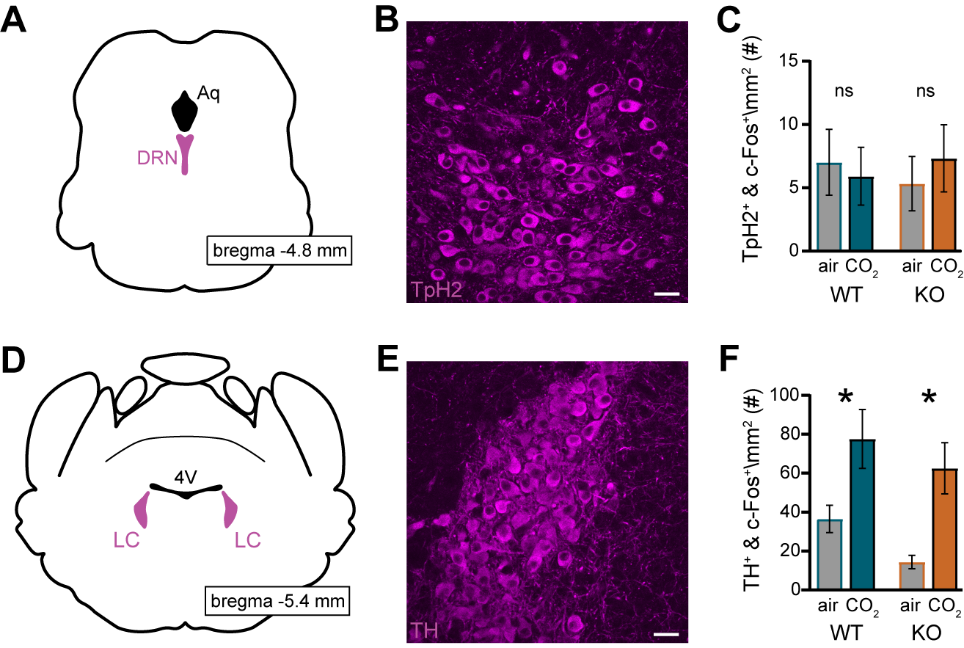

**Supplementary Fig S2. Serotonin and noradrenaline neuron activity following CO_2_ inhalation is the same in GluD1-WT and GluD1-KO mice.**

**A** Schematic of the mouse coronal brain slice highlighting the location of the dorsal raphe nucleus (DRN). **B** Representative maximum intensity projection confocal image of serotonin neurons in the DRN, identified via immunostaining for tryptophan hydroxylase 2 (TpH2); scale bar, 25 µm. **C** Quantification of TpH2^+^ and c-Fos^+^ neurons in the DRN from GluD1-WT (blue) and GluD1-KO (orange) mice (WT, p=0.763, n=9-10; KO, p=0.550, n=11-12, 2-way ANOVA with Fischer’s LSD test). **D** Schematic of the mouse coronal brain slice highlighting the location of the locus coeruleus (LC). **E** Representative maximum intensity projection confocal image of noradrenaline neurons in the LC, identified via immunostaining for tyrosine hydroxylase (TH); scale bar, 25 µm. **F** Quantification of TH^+^ and c-Fos^+^ neurons in the LC from GluD1-WT (blue) and GluD1-KO (orange) mice (WT, p=0.016, n=9-10; KO, p=0.003, n=11-12, 2-way ANOVA with Fischer’s LSD test).

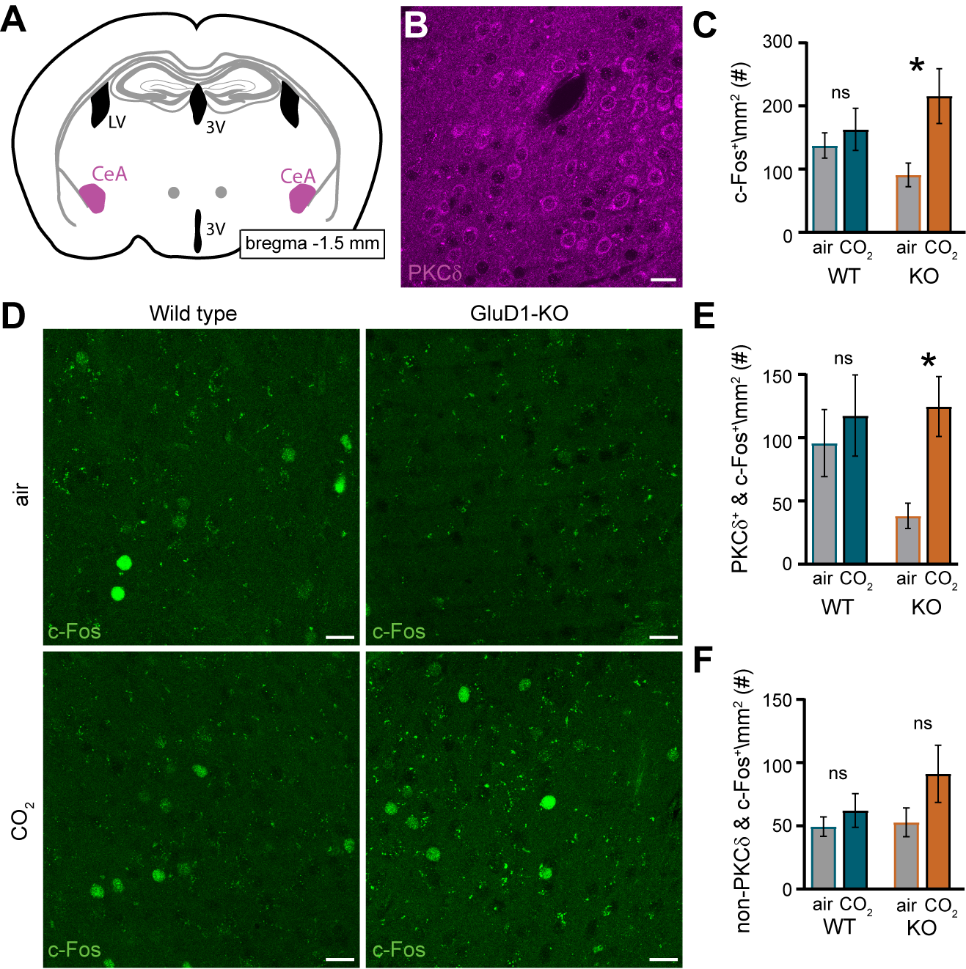

**Supplementary Fig S3**. **PKCδ^+^ neuron activity in the CeA is increased following elevated CO_2_ exposure in GluD1-KO mice.**

**A** Schematic of the mouse coronal brain slice highlighting the location of the central amygdala (CeA). **B** Representative maximum intensity projection confocal image of PKCδ neurons in CeA; scale bar, 20 µm. **C** Quantification of c-Fos^+^ expression in the CeA from GluD1-WT (blue) and GluD1-KO (orange) mice (WT, p=0.584, n=10; KO, p=0.005, n=12, 2-way ANOVA with Fischer’s LSD test). **D** Representative maximum intensity projection confocal images of c-Fos expression in the CeA from GluD1-WT (left column) and GluD1-KO (right column) mice following exposure to room air (top row) or CO_2_ (bottom row); scale bars, 20 µm. **E** Quantification of PKCδ^+^ and c-Fos^+^ neurons in the CeA from GluD1-WT (blue) and GluD1-KO (orange) mice (WT, p=0.586, n=10; KO, p=0.005, n=12, 2-way ANOVA with Fischer’s LSD test). **F** Quantification of non-PKCδ^+^ and c-Fos^+^ neurons in the CeA from GluD1-WT (blue) and GluD1-KO mice (WT, p=0.680, n=10; KO, p=0.090, n=12, 2-way ANOVA with Fischer’s LSD test).
